# Efficiency of neutral honey as a tissue fixative in histopathology

**DOI:** 10.1101/2021.04.27.437988

**Authors:** Nasar Alwahaibi, Buthaina Al Dhahli, Halima Al Issaei, Loai Al Wahaibi, Shadia Al Sinawi

**Affiliations:** Department of Allied Health Sciences, College of Medicine and Health Sciences, Sultan Qaboos University. Oman; Department of Pathology, Sultan Qaboos University Hospital, Sultan Qaboos University. Oman

**Keywords:** Histopathology, fixation, honey, formalin, neutral buffered

## Abstract

In the routine laboratory, 10% neutral buffered formalin (NBF) is the fixative of choice. However, formalin is a human carcinogen. To the best of our knowledge, neutral honey, not natural or artificial honey, has not been tested to fix histological tissues. This study aimed to examine the efficiency of neutral buffered honey and other types of honey fixatives to fix histological tissues. The most two natural common Omani honey were used as fixatives, namely Sumar and date. We tested samples of rat liver, kidney, and stomach. Nine types of fixatives were used. All tissues were treated equally. The evaluation was performed blindly by three senior biomedical scientists who work in a histopathology laboratory. Hematoxylin and eosin showed adequate staining in all groups when compared to 10% NBF. The intensity and specificity of Jones Methenamine silver stain in 10% Sumer and Date honey and 10% alcoholic Sumer honey showed similar findings of 10% NBF. The specificity and intensity of all groups for Periodic acid–Schiff were comparable with 10% neutral buffered formalin accepts for 10% Sumer honey and 10% Alcoholic Date honey. However, all honey groups showed weak staining for the reticulin fibers using Gordon and Sweets method. Vimentin showed comparable findings with 10% NBF as there were no significant differences. The findings of this study are promising. Further in depth research on honey as a possible safe substitute fixative for formalin should be conducted.

## Introduction

In histopathology, fixation is an initial and the most critical step in the tissue processing which is important for good microscopic examination.^1^ The purpose of the fixation is to preserve the tissue as in life-like condition and this is achieved by preventing autolysis and bacterial putrefaction.^2^ In routine laboratory, 10% neutral buffered formalin is the fixative of choice because it is ready available, has international acceptance, its preparation requires less time, cheap, fast technique, good prolonged long-term storage, suitable for post fixation processing and easy to use as well as its ability to prevent autolysis and bacterial growth.^3,4^ However, the International Agency for Research on Cancer (IARC) and U.S. Environmental Protection Agency (EPA) classified the formaldehyde, which is the basic component of formalin, as a human carcinogen.^5^ In addition, formalin can affect skin and mucous membrane causing allergic contact dermatitis and irritation of the mucous membranes of the mouth and upper respiratory tract and it can affect respiratory tract causing coughing, sneezing, laryngospasm, temporary reversible decrease in lung function, pulmonary edema, degenerative diseases, inflammatory and hyperplastic changes of the nasal mucosa, chronic rhinitis, asthma and many more.^6^ Also, the extraction of intact nucleic acids (DNA and mRNA) for molecular tests, from samples processed by formalin is problematic or impossible due to a DNA-protein or DNA-DNA crosslinks induced by formalin.^7^

Therefore, formalin should be replaced by a safer alternative fixative such as natural fixatives which are eco-friendly, economical and ready available substances. Example of natural fixatives are honey, sugar, jaggery, molasses, saline, rose water, coconut oil and others. Honey is a mixture of sugars and other compounds such as several minerals, trace elements, vitamins such as vitamin C, ascorbic acid, hydrogen peroxide, pinobanksin, pinocembrin, chrysin and catalase.^7^. These compounds give distinct properties to the honey which are antiautolytic, antimicrobial, antiviral, antimutagenic and antioxidant effects which have been known for several centuries.^4^ Honey also has been shown to possess acidic, preserving, dehydrating and tissue hardening properties. In addition, it can penetrate the deepest tissue.^3^ These properties make the honey as a potential tissue fixative in terms of fixation. Previous studies showed that tissues fixed in honey at low pH were less firm after fixation, homogenization of connective tissue, breach in continuity of sections, folding of the tissue sections and growth of molds over a period of time.^1,3,6,7^ In addition, higher concentrations were found to cause tissue shrinkage and loss of tissue architecture due to the high osmolality of honey. To the best of our knowledge, neutral honey has not been experimented to fix histological tissues. Thus, we aimed to examine the efficiency of neutral buffered honey to fix histological tissues.

## Methods

We tested samples of rat liver, kidney and stomach. Three pieces from each organ were used. All tissues were treated similarly. In order to have a uniform thickness (3 mm) for all groups, tissues were cut using the Cutmate forceps (Forceps for 2/3/4 mm tissue blocks, Code No 62359, Milestone s.r.l, Sorisole, Italy). The most two common natural Omani honeys were used as fixatives, namely Sumer and date.

The tissues were immediately placed in fixatives as shown in Table 1. In each group, formalin was used as the gold standard. The pH of all fixatives were detected using pH meter (pH meter, Extech® Instrument, WQ510, Taiwan). For neutral buffers (Sumer and Date honeys and formalin) sodium phosphate monobasic and sodium phosphate dibasic were used. After 24 hours’ fixation at room temperature (18 – 23 °C), all tissues were processed as previously described using an automated histoprocessor (Spin Tissue Processor Microm STP 120; Thermo Scientific, Walldorf, Germany).^8^ In all tissues, 3 μm sections were cut using a rotatory microtome (Leica RM2135, Nussloch, Germany). Nine slides from each organ were cut.

**Table 1:** Experimental design.

|  | 10% Neutral buffered |  |  | 10% fixatives |  |  | 10% alcoholic fixatives |  |  |
| --- | --- | --- | --- | --- | --- | --- | --- | --- | --- |
| Groups | NBF | NBS | NBD | F | S | D | AF | AS | AD |
| pH | 7.20 | 7.25 | 7.27 | 3.62 | 3.56 | 5.15 | 3.95 | 5.10 | 5.85 |
| Liver | 3 | 3 | 3 | 3 | 3 | 3 | 3 | 3 | 3 |
| Kidney | 3 | 3 | 3 | 3 | 3 | 3 | 3 | 3 | 3 |
| Stomach | 3 | 3 | 3 | 3 | 3 | 3 | 3 | 3 | 3 |

NBF: Neutral buffered formalin. NBS: Neutral buffered Sumer honey. NBD: Neutral buffered Date honey. F: Formalin. S: Sumer honey. D: Date honey. AF: Alcoholic formalin. AS: Alcoholic Sumer honey. AD: Alcoholic Date honey. 3: number of pieces.

All slides were stained with hematoxylin and eosin (H & E).^8^ In addition, a number of special stains were used to further evaluate the efficacy of honeys in different groups.^8^ Jones’ Methenamine silver stain (JMS) to stain glomerular basement membranes in kidney. Gordon and Sweets (G&S) method to stain reticular fibers. Periodic acid–Schiff (PAS) method to detect basement membranes of glomerular capillary loops and tubular epithelium. All special stains were performed using the Ventana BenchMark Special Stains system (Ventana Medical Systems, Inc., Code No: 06657389001, Tucson, AZ, USA). For immunohistochemistry, all tissue blocks were cut at 4 μm using a rotatory microtome (Leica RM2135; Nussloch, Germany). Slides were deparaffinized, rehydrated and epitope retrieved by PT link (Code PT200, Agilent Dako, CA, USA). All slides were then washed three times in phosphate buffered saline (PBS) each for 5 min. After that, slides were incubated with primary vimentin antibody (ab137321, Abcam plc, Cambridge, U.K.) at 1:1000 dilution for 30 minutes. Slides were then washed with PBS three times each for 5 min followed by incubation for 30 min with secondary antibody (Envision Flex, High pH (Link), Hrp. Rabbit/Mouse. Code No K8000, Dako, CA, USA) and then washed with PBS three times each for 5 min. After that, the reaction was visualized using diaminobenzidine (Code No K3468, Dako, CA, USA) for 2 min. Slides were counterstained with Mayer’s hematoxylin for 2 min and then washed for 2 min in running tap water. Finally, slides were dehydrated, cleared and mounted in dibutylphthalate polystyrene xylene.

## Evaluation and data analysis

All slides were examined by light microscopy (BX40, Olympus Optical Co, Tokyo, Japan) attached with camera (DP71 controller, Olympus Optical Co, Tokyo, Japan). The cost of formalin, Sumer and date honeys were obtained from local suppliers. For H& E stain, nuclear and cytoplasmic staining, cell morphology, clarity of staining and uniformity of staining criteria were used to assess the quality of all sections.^9^ A score of 0 and 1 was given to each parameter and then scores were added up to be graded. If the score was ≤ 2, graded as inadequate and if the score was 3-5, graded as adequate.^10^ For special stains, specificity and intensity were used and graded either negative (0), weak (1), moderate (2) or strong (3). For immunohistochemistry, intensity and background were used and graded either negative (0), weak (1), moderate (2) or strong (3). Evaluation was performed blindly by three senior biomedical scientists who work in histopathology laboratory. The data were analyzed using the Statistical Package for the Social Sciences (SPSS) software version 23 (SPSS Inc., Chicago, IL, USA). Fisher’s exact and ANOVA tests were used to compare the two measurements of the variable. A *P* value of less than 0.05 was considered statistically significant.

## Results

In all groups, there were no noticeable differences in sectioning, formation of ribbons, and floating on the water bath. According to the nuclear and uniformity of staining, there was no significant difference among all the groups. By contrast, 10% neutral buffered Sumer honey, 10% neutral buffered Date honey, 10% Sumer honey and 10% Date honey showed significant difference with regard to cytoplasmic staining (Figure 1). Other staining criteria are shown in Table 2.

**Figure 1:**
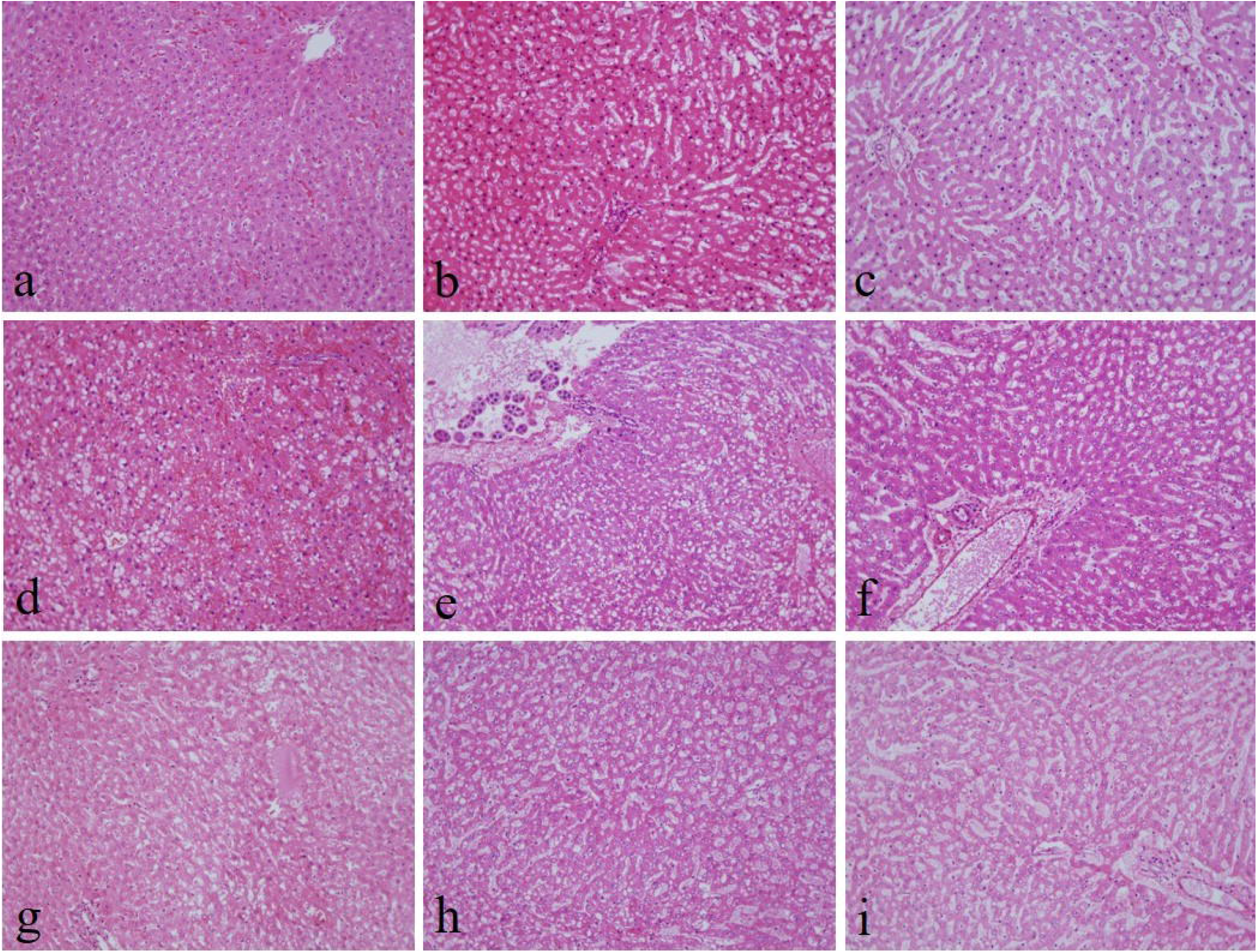
Hematoxylin and eosin stained liver sections of (a) 10% neutral buffered formalin, (b) 10% neutral buffered Sumer honey, (c) 10% neutral buffered Date honey, (d) 10% formalin, (e) 10% Sumer honey, (f) 10% Date honey, (g) 10% alcoholic formalin, (h) 10% alcoholic Sumer honey and (i) 10% alcoholic Date honey (x200).

**Table 2:** Comparison of different honey fixative groups using hematoxylin and eosin staining method.

|  | 10% Neutral buffered |  |  | 10% fixatives |  |  | 10% alcoholic fixatives |  |  |
| --- | --- | --- | --- | --- | --- | --- | --- | --- | --- |
| Groups | NBF | NBS | NBD | F | S | D | AF | AS | AD |
| Nuclear Staining |  |  |  |  |  |  |  |  |  |
| Adequate | 9 | 6 | 6 | 9 | 8 | 8 | 9 | 9 | 9 |
| Inadequate | 0 | 3 | 3 | 0 | 1 | 1 | 0 | 0 | 0 |
| *P* value |  | 0.206 | 0.206 | 1.00 | 1.00 | 1.00 | 1.00 | 1.00 | 1.00 |
| Cytoplasm Staining |  |  |  |  |  |  |  |  |  |
| Adequate | 9 | 0 | 3 | 9 | 3 | 3 | 9 | 9 | 6 |
| Inadequate | 0 | 9 | 6 | 0 | 6 | 6 | 0 | 0 | 3 |
| *P* value |  | 0.000 | 0.009 | 1.00 | 0.009 | 0.009 | 1.00 | 1.00 | 0.206 |
| Cell Morphology |  |  |  |  |  |  |  |  |  |
| Preserved | 9 | 5 | 6 | 9 | 8 | 3 | 9 | 3 | 0 |
| Unpreserved | 0 | 4 | 3 | 0 | 1 | 6 | 0 | 6 | 9 |
| *P* value |  | 0.082 | 0.206 | 1.00 | 1.00 | 0.009 | 1.00 | 0.009 | 0.000 |
| Clarity of staining |  |  |  |  |  |  |  |  |  |
| Present | 9 | 0 | 3 | 9 | 6 | 9 | 9 | 6 | 6 |
| Not present | 0 | 9 | 6 | 0 | 3 | 0 | 0 | 3 | 3 |
| *P* value |  | 0.000 | 0.009 | 1.00 | 0.206 | 1.00 | 1.00 | 0.206 | 0.206 |
| Uniformity of staining |  |  |  |  |  |  |  |  |  |
| Present | 9 | 6 | 6 | 9 | 9 | 6 | 9 | 9 | 9 |
| Not present | 0 | 3 | 3 | 0 | 0 | 3 | 0 | 0 | 0 |
| *P* value |  | 0.206 | 0.206 | 1.00 | 1.00 | 0.206 | 1.00 | 1.00 | 1.00 |

NBF: neutral buffered formalin. NBS: neutral buffered Sumer honey. NBD: neutral buffered Date honey. F: formalin. S: Sumer honey. D: Date honey. AF: Alcoholic formalin. AS: Alcoholic Sumer honey. AD: Alcoholic Date honey.

The intensity and specificity of JMS in 10% Sumer and Date honeys and 10% alcoholic Sumer honey showed similar findings of 10% NBF (Figure 2). In addition, the specificity and intensity of all groups for PAS were comparable with 10% neutral buffered formalin accepts for 10% Sumer honey and 10% Alcoholic Date honey were the intensity was significant in comparison with the 10% neutral buffered formalin (Figure 3). However, all honey groups showed week staining of the reticulin fibers using G&S method (Figure 4) (Table 3). Immunohistochemical staining with vimentin, showed comparable findings with 10% NBF as there were no significant differences noted for all groups (Table 4) (Figure 5). Sumer honey is more expensive than formalin. One liter of Sumer costs 60 Omani Rial (OMR) which is equivalent to 155.79 USD whereas formalin costs 1.6 OMR equivalent to 4.15 USD. Date honey costs 5.19 USA.

**Figure 2:**
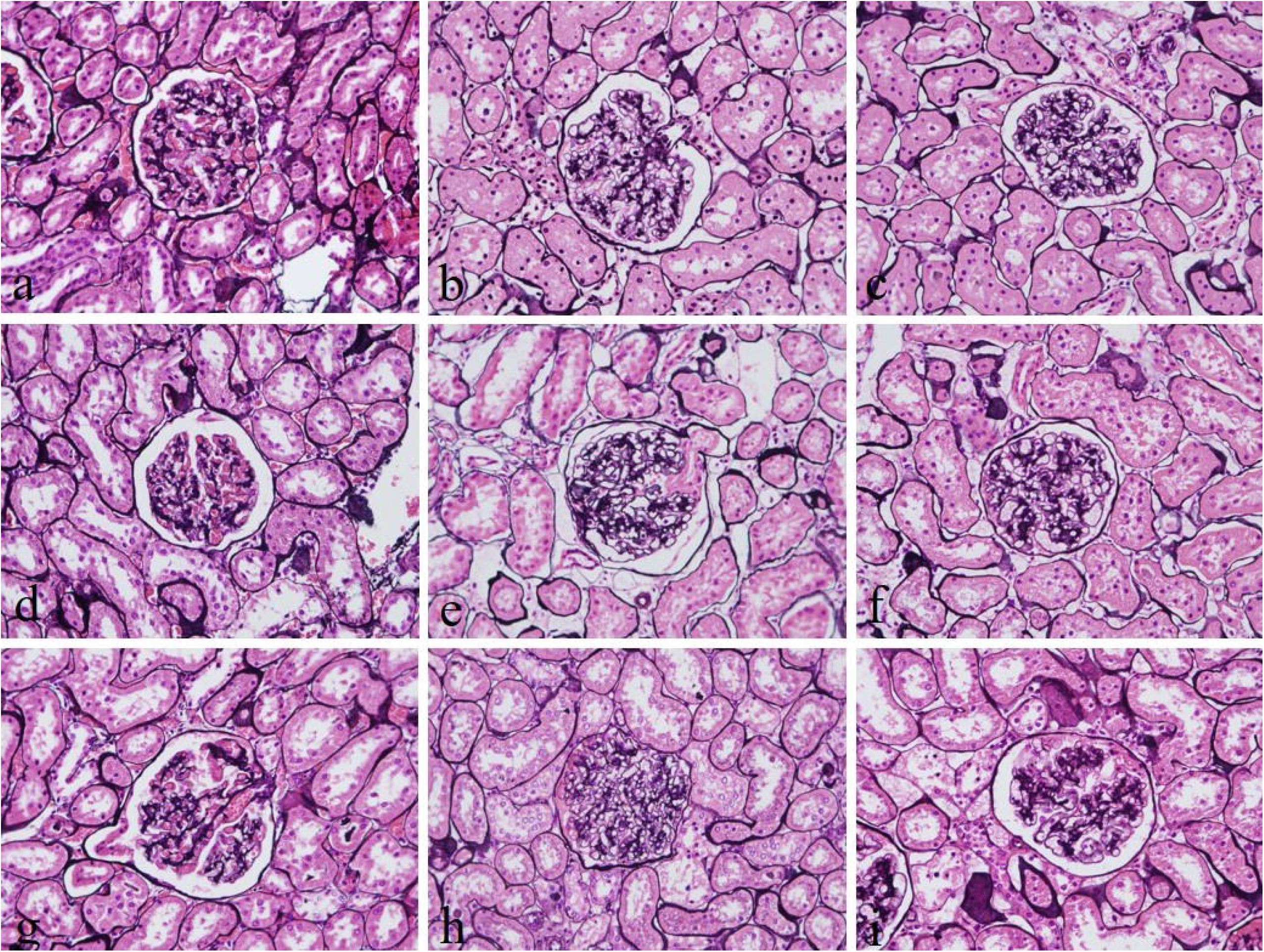
Jones’ Methenamine silver stain of (a) 10% neutral buffered formalin, (b) 10% neutral buffered Sumer honey, (c) 10% neutral buffered Date honey, (d) 10% formalin, (e) 10% Sumer honey, (f) 10% Date honey, (g) 10% alcoholic formalin, (h) 10% alcoholic Sumer honey and (i) 10% alcoholic Date honey (x400).

**Figure 3:**
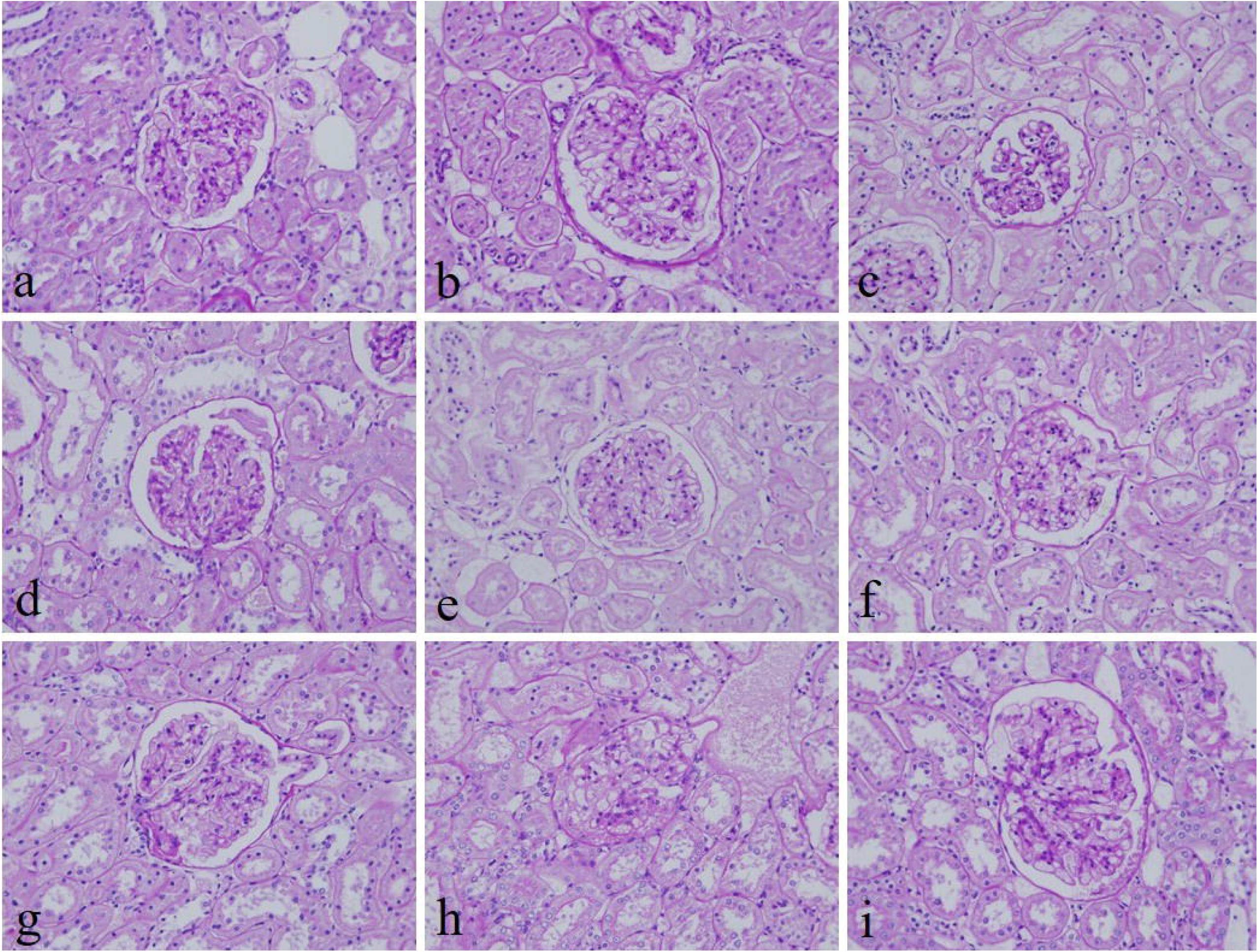
Periodic acid–Schiff stain of (a) 10% neutral buffered formalin, (b) 10% neutral buffered Sumer honey, (c) 10% neutral buffered Date honey, (d) 10% formalin, (e) 10% Sumer honey, (f) 10% Date honey, (g) 10% alcoholic formalin, (h) 10% alcoholic Sumer honey and (i) 10% alcoholic Date honey (x400).

**Figure 4:**
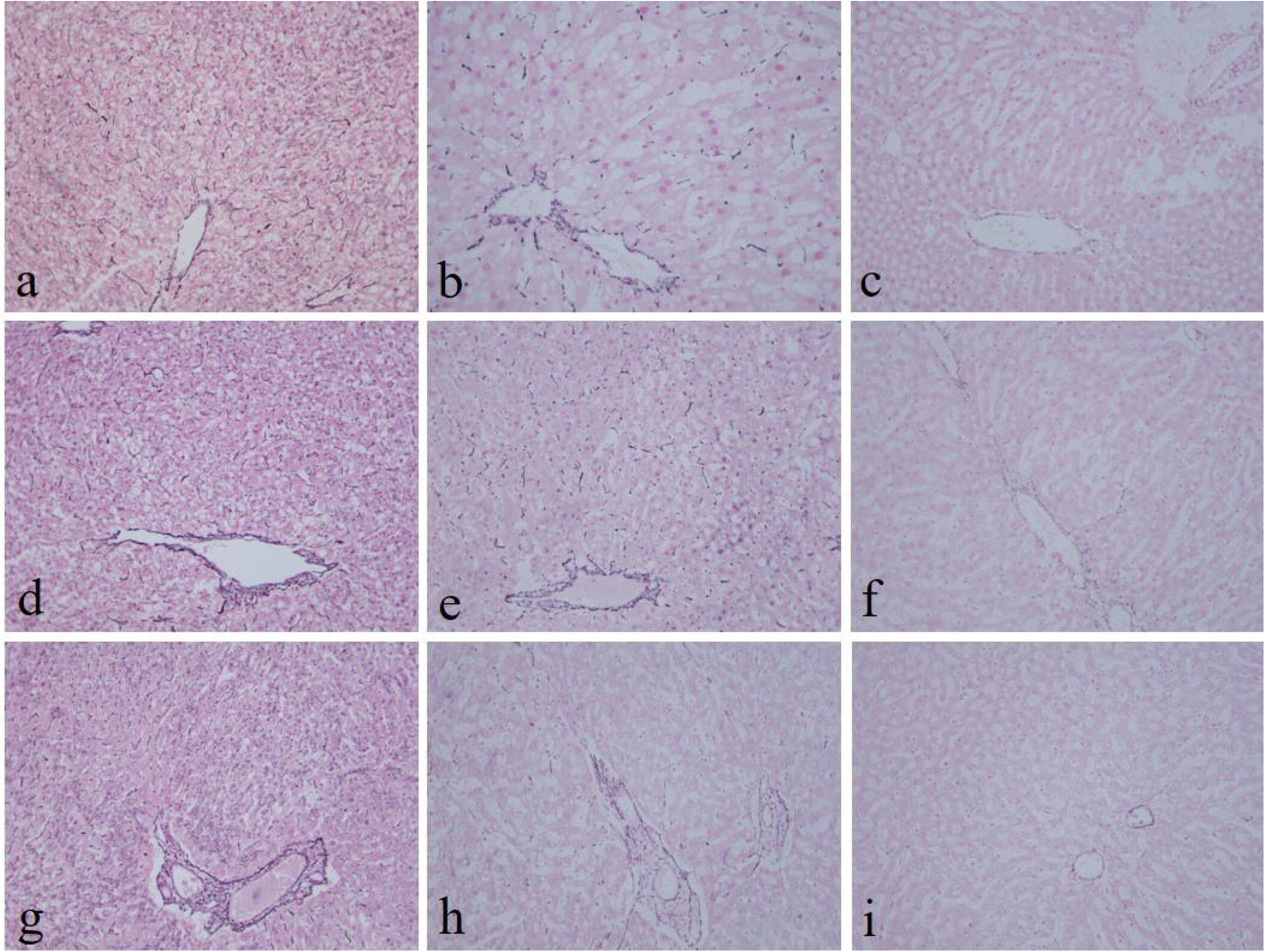
Gordon and Sweets stain of (a) 10% neutral buffered formalin, (b) 10% neutral buffered Sumer honey, (c) 10% neutral buffered Date honey, (d) 10% formalin, (e) 10% Sumer honey, (f) 10% Date honey, (g) 10% alcoholic formalin, (h) 10% alcoholic Sumer honey and (i) 10% alcoholic Date honey (x400).

**Table 3:**
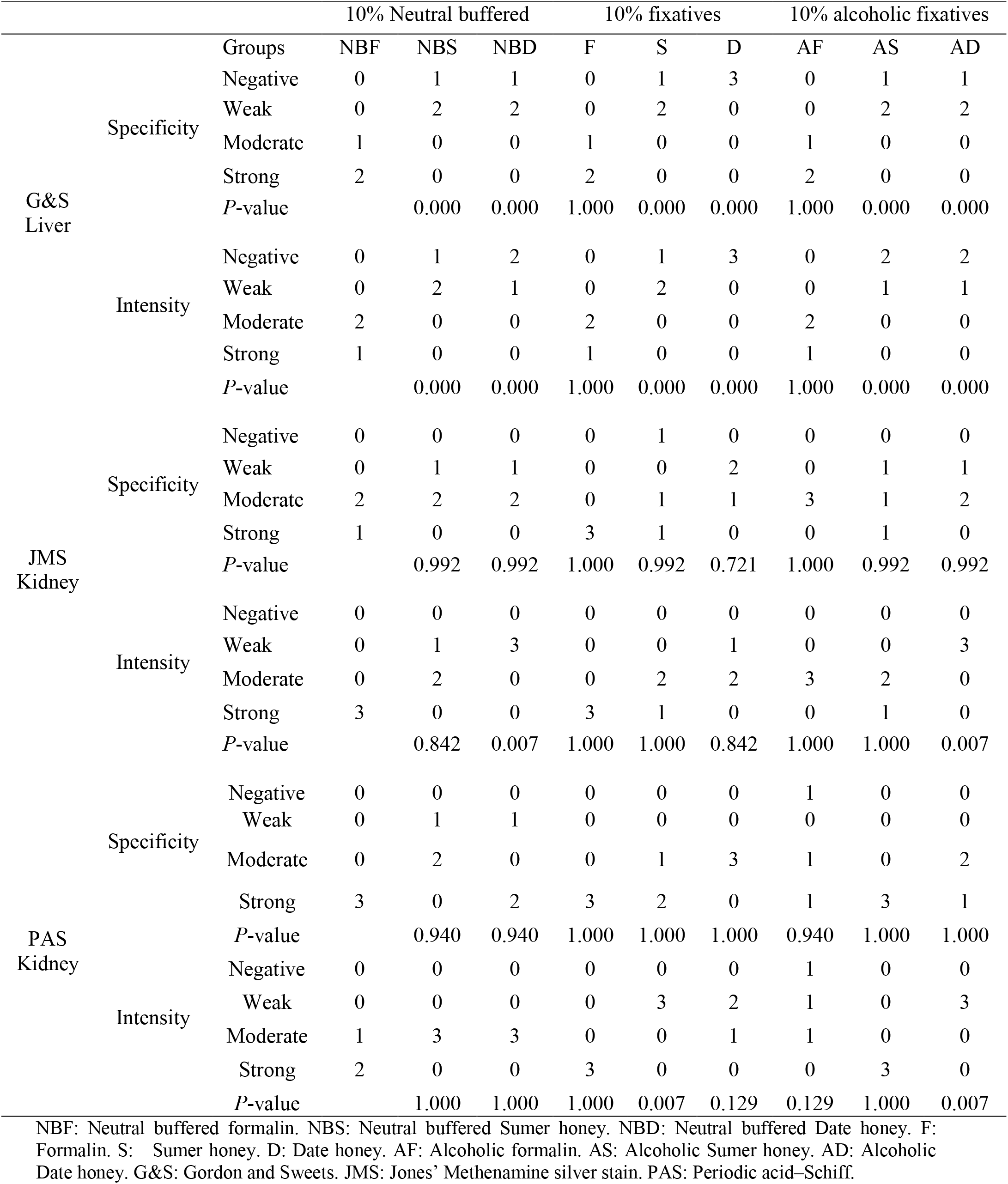
Comparison of different honey fixative groups using three special stains.

NBF: Neutral buffered formalin. NBS: Neutral buffered Sumer honey. NBD: Neutral buffered Date honey. F: Formalin. S: Sumer honey. D: Date honey. AF: Alcoholic formalin. AS: Alcoholic Sumer honey. AD: Alcoholic Date honey. G&S: Gordon and Sweets. JMS: Jones’ Methenamine silver stain. PAS: Periodic acid–Schiff.

**Table 4:**
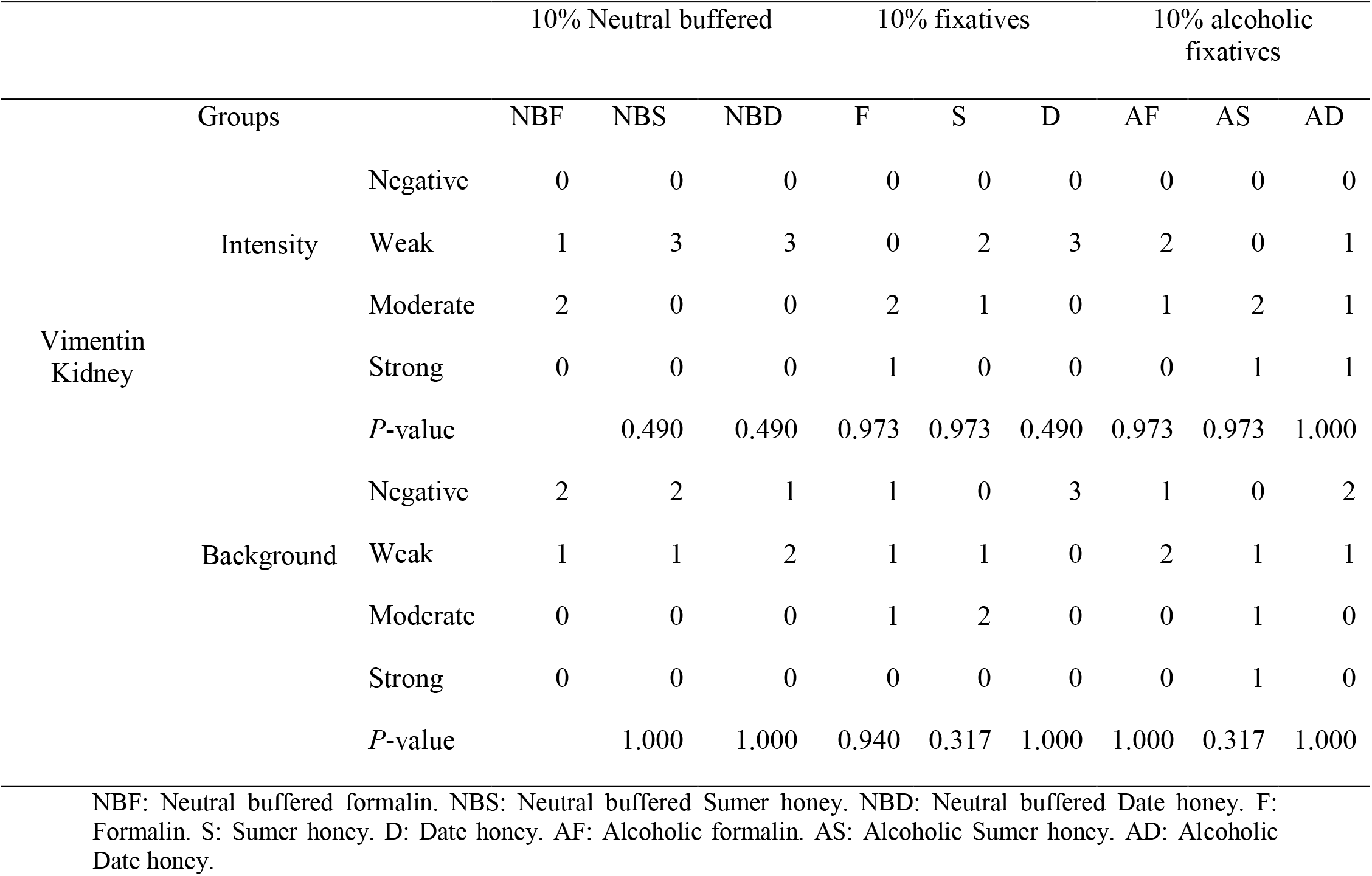
Comparison of different honey fixative groups using vimentin marker.

NBF: Neutral buffered formalin. NBS: Neutral buffered Sumer honey. NBD: Neutral buffered Date honey. F: Formalin. S: Sumer honey. D: Date honey. AF: Alcoholic formalin. AS: Alcoholic Sumer honey. AD: Alcoholic Date honey.

**Figure 5:**
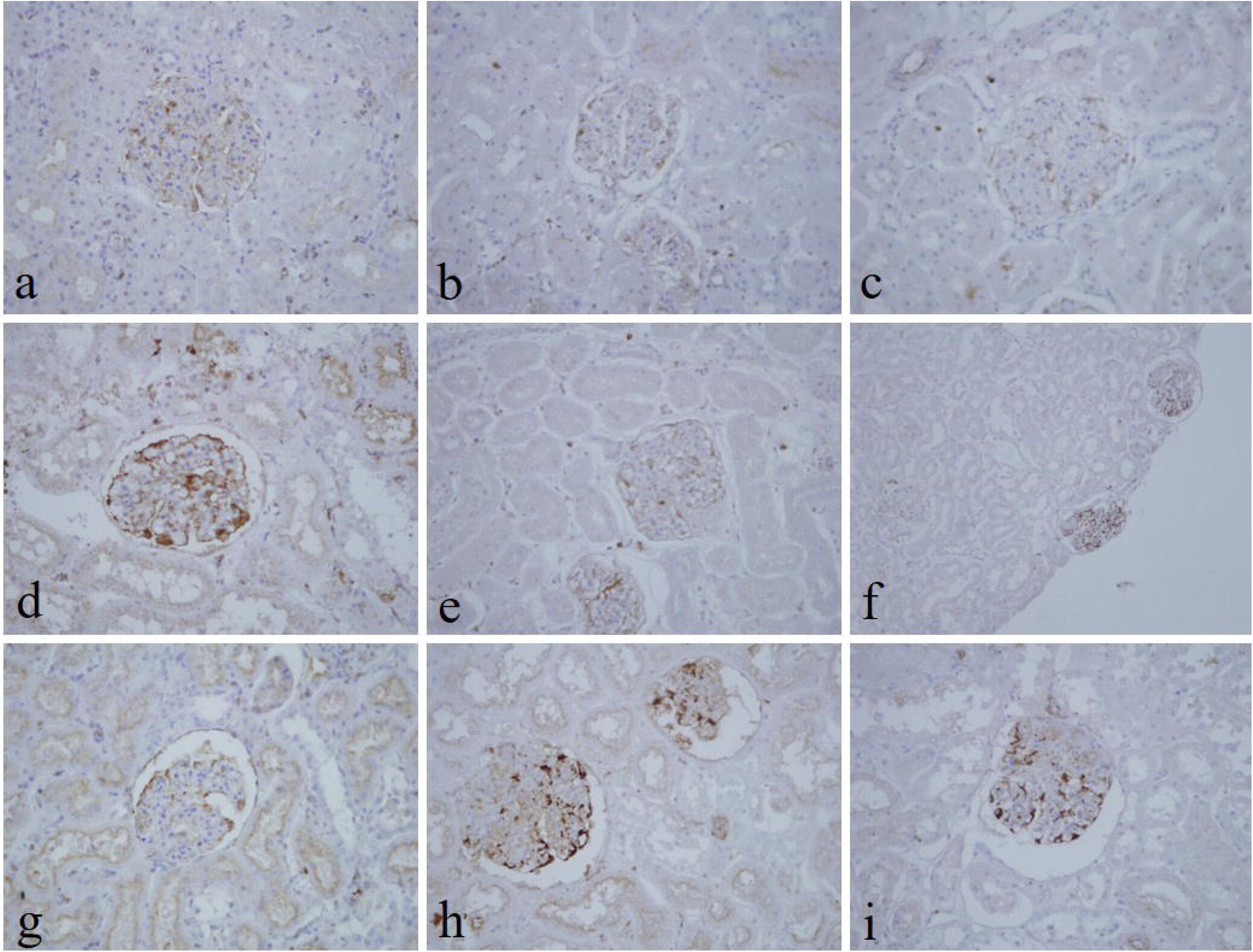
Immunohistochemical staining for vimentin in kidney sections fixed in (a) 10% neutral buffered formalin, (b) 10% neutral buffered Sumer honey, (c) 10% neutral buffered Date honey, (d) 10% formalin, (e) 10% Sumer honey, (f) 10% Date honey, (g) 10% alcoholic formalin, (h) 10% alcoholic Sumer honey and (i) 10% alcoholic Date honey (x400).

## Discussion

To the best of our knowledge, this is the first study to evaluate honey as a neutral buffered honey similar to that of the neutral buffered formalin. In this study, we evaluated two types of honey with different formula and compared with the gold standard fixative which is 10% neutral buffered formalin. In this study, two types of honey were used, Al Baram (Sumar) and date. Al Baram is the most well-known and one of the finest types of Omani natural honey. It is a light black in colour and produced from Omani Dwarf bees. It is very expensive honey as it is produced in small amounts and usually found in the caves of the mountains, therefore, it is difficult to acquire. Date honey is very common in Oman as it is extracted from the dates. It is slightly brown in colour and not expensive.

Literature has revealed two possible mechanisms for honey fixation. The first one is through the conversion of carbohydrate to gluconic acid, which is known to have a wide application in food and pharmaceutical industry as a preservative by preventing the decomposition of the food. The other one is through the presence of fructose in honey which at low pH breaks down to form aldehyde groups. Subsequently, these aldehyde groups form methylene bridges with lysine amino acids, which lead to the tissue fixation. This mechanism of honey fixation is similar to the action of formaldehyde.^11–14^ In this study, we have chosen experimental conditions such as a temperature of 18 – 23 °C, 24 hours’ fixation time, tissue thickness of 3 mm and fixation volume of 1:10. In line with other study, previous study revealed 100% penetration and 70% binding rates at a temperature of 20 – 22 °C using 3 mm thick tissue slices and for 24 hours’ fixation.^15^ In fact, 1:10 fixation volume has been reported by many researchers and found to be the most recommended ratio.^15,16^ In addition, during our pilot study we used 5%, 10%, 15% honey fixative concentrations for 24 and 48 hours’ fixation time and found that 10% for 24 hours’ fixation preserved tissue morphology well.

The present study evaluates three important aspects of honey fixation: hematoxylin and eosin (H&E), special stain and immunohistochemistry. In both types of honeys and in all groups, H & E results showed that overall quality of tissue staining is comparable with the 10% neutral buffered formalin. The nucleus was well preserved with clear nuclear details. It is known that honey contains ascorbic acid as well as various vitamins, carbohydrates, minerals and trace elements. Thus, it makes honey at low pH. Low pH fixative is good for nuclear staining.^17^

The finding of the present study is in line with other study which reported that 10% honey, using rat liver and kidney tissues, at room temperature for 24-hour fixation gave comparable results with those obtained by formalin-fixed control tissues.^16^ Another similar study which used the same staining criteria to evaluate honey as a substitute for formalin found that 10% honey is as good as 10% neutral buffered formalin and suggested that honey is a safe alternative for formalin.^6^ Recent two studies in cytology showed that 20% honey fixed oral smears had acceptable nuclear and cytoplasmic staining, well preserved cell morphology, clarity and uniformity of staining as comparable to ethanol, as a gold standard fixative in cytology, with no statistical difference between both fixatives.^18,19^ However, in the current study, cytoplasmic staining was inadequate in neutral buffered Sumer honey, neutral buffered Date honey, 10% Sumer honey and 10% Date honey. We thought by increasing the pH in neutral buffered Sumer honey and neutral buffered Date honey fixatives would make the cytoplasmic staining enhanced. The results of the present study are in concordance with other similar study where they compared formalin fixed tissues with honey fixed tissues, and concluded that the nuclear details are well established compared to the cytoplasm. ^7^ Very recent study showed 10% alcoholic honey turned out to be a good fixative with formalin-like effects, however 10% honey solution is not a suitable fixative for preserving large tissue samples.^20^

Reticulin fibers in all groups were not well demonstrated. The staining was week. This finding disagrees with other study which reported that reticulin fibers using silver impregnation were well demonstrated using 10% honey as a fixative. ^21^ However, the demonstration of glomerular basement membranes in kidney by JMS in 10% Sumer and Date honeys and 10% alcoholic Sumer honey showed similar findings of 10% NBF. Another study used different specials stains to assess the efficacy of 10% honey as a fixative agent for the preservation of cellular and structural characteristics. They found that Masson’s trichrome and Van Gieson staining results are similar to those fixed by formalin.^22^ Similarly, the specificity and staining intensity using PAS on honey fixed tissues were comparable with 10% neutral buffered formalin.^4^ our results are in line with this study where PAS stain in all groups revealed similar findings with 10% neutral buffered formalin. However, the intensity for only 10% Sumer honey and 10% Alcoholic Date honey were significant in comparison with the 10% neutral buffered formalin.

In the current study, both honeys in H & E and special stains showed the absence of red blood cells in liver, kidney and stomach tissues. When red blood cells are exposed to hydrogen peroxide, which is a component of honey, this would make cellular changes that lead to alterations in phospholipid organization and cell shape and membrane deformability. Hydrogen peroxide has the ability to form covalent complex with hemoglobin and spectrin which are specific structures of red blood cells.^23^ This may explain why honey masks the staining of red blood cells. We assumed that alcoholic honey would stain red blood cells as alcohol is a good fixative for red blood cells but it did not work.

In the current study, vimentin as an immunohistochemical marker was evaluated. In routine immunohistochemistry, this marker is used as an internal control for detecting cytoplasmic staining.^24,25^ Vimentin in all honey groups showed similar findings of 10% NBF. The finding of the current study is in line with other similar study where they reported that honey fixation was similar to neutral buffered formalin fixation in demonstrating ki-67 and vimentin markers in different fresh tissues including endometrium, breast, placenta, uterus, omentum, suprarenal, stomach and lung.^2^ Another study reported that good levels of staining without antigen retrieval for common leukocyte antigen, cytokeratin AE1/AE3 and epithelial membrane antigen in breast tumor samples treated with hone.^26^ In addition, the demonstration of vimentin and pan-cytokeratin in gingiva tissues using 10% honey was similar with formalin fixed tissues.^22^ Similarly, the immunhistochemical demonstration of pan-cytokeratin was comparable using 20% honey fixative in fresh goat oral mucosa.^27^

The findings of this study encourage pharmaceutical companies to produce high quality commercial honeys to be experimented for possible tissue fixation. For example, 10% pure honey, 10% neutral buffered honey and 10% alcoholic honey. However, further future study is highly recommended. It is important to note that the cost of honey is more than formalin, in particular, with Sumer honey. However, the safety of workers is more important.

Several limitations of our study are worth noting. First, we should point out that large tissues or small biopsies were not evaluated, different sized tissues would produce more meaningful results. Second, only one immunohistochemical marker was evaluated, a wide range of immunohistochemical markers would give a better picture of how good or bad the honey fixative. Third, limited number of special stains were evaluated. Finally, DNA/RNA extraction from honey fixed specimens were not assessed.

In conclusions, the findings of this study are promising and encouraging. Further in-depth research on honey as a possible safe substitute fixative for formalin should be conducted.

## Notes

### Competing Interest Statement

The authors have declared no competing interest.

